# Characterization of novel fungal-algal symbiosis on LDPE plastic surfaces in the Mediterranean Sea

**DOI:** 10.1101/2023.06.17.545398

**Authors:** Sheli Itzahri, Keren Davidov, Matan Oren

## Abstract

Plastic debris in the ocean serves as a stable ground for the formation of a complex ecosystem, termed plastisphere, which includes a variety of organisms from different taxonomic groups. Not much is known about the relationships between the organisms of the plastisphere communities. In this study we describe a novel symbiotic relationship between a marine fungus and several species of diatoms on plastic surfaces that were submerged in the water of a Mediterranean Sea marina in Israel. Scanning electron microscope images of the surfaces revealed a network of fungal hyphae with multiple diatom cells attached to them via the side or the tip of their body. Using DNA metabarcoding for the fungal Internal Transcribed Spacer (ITS) barcode locus, we found that the symbiotic fungus belongs to the phylum Ascomycota, and that it is more abundant on low density polyethylene (LDPE) surfaces compared to other plastic polymers and glass. We hypothesize that the observed symbiotic relationship may have mutual benefits for both parties, including surface-anchoring for the diatoms and nutritional benefits for the fungus, that reflects a recent adaptation for life on floating plastic debris.

## Introduction

Marine microplastic debris is now more abundant than ever before with a recent estimate of 82–358 trillion particles floating in the world oceans (Eriksen, Cowger et al. 2023). The plastic surfaces serve as an ideal substrate for the colonization of complex microbial communities forming a man-made ecosystem - the plastisphere, which is distinguished from that of its surrounding water (Amaral-Zettler, Zettler et al. 2020). Most microplastic debris in the oceans floats near the surface and benefits from high exposure to sunlight, allowing the establishment of complex communities that include prokaryotic, and eukaryotic phototrophs alongside primary consumers, predators and saprotrophs (Amaral-Zettler, Zettler et al. 2020). Some of the most abundant microorganisms within the plastisphere of floating plastic debris are diatoms and the fungi (Amaral-Zettler, Zettler et al. 2020, Du, Liu et al. 2022).

The diatoms are unicellular algae that are enclosed in a silica skeleton termed ‘frustule’ which is made of two overlapping valves. The detailed structures of the frustules as observed by Scanning Electron Microscope (SEM), are often used as the basis for high-resolution taxonomical classification (Round, Crawford et al. 1990). Diatoms reproduce asexually through binary fission as well as by sexual reproduction following meiosis (Vyverman 2004). Overall, the diatoms are estimated to carry out one-fifth of the earth’s photosynthesis (Armbrust 2009) and have a major role in the ocean carbon and silica cycles (Benoiston, Ibarbalz et al. 2017). Diatoms have been consistently found in high abundance within the plastisphere (Carson, Nerheim et al. 2013, Zettler, Mincer et al. 2013, Reisser, Shaw et al. 2014, Davidov, Iankelevich-Kounio et al. 2020) and are amongst the first to colonize plastic surfaces (Amaral□Zettler 2022).

Using DNA metabarcoding, few studies showed the presence of fungi on plastic surfaces either collected from the marine environment (Lacerda, Proietti et al. 2020, Gkoutselis, Rohrbach et al. 2021, Kim, Lee et al. 2022) or from surfaces that were immersed in it (Davidov, Iankelevich-Kounio et al. 2020, Marsay, Ambrosino et al. 2023, Philippe, Noël et al. 2023). We have previously shown that plastic surfaces that were incubated for one month in the Herzelia marina, Israel, are covered with a biofilm that includes a rich fungal community and often a network of fungal hyphae (Marsay, Koucherov et al. 2022). In this study, we describe a novel type of symbiotic relationship between a fungus and several species of diatoms that took place on low-density polyethylene (LDPE) plastic surfaces.

## Methods

### Experiment setup and sampling

LDPE bags (20 cm x 18 cm) were incubated for one month in the water (20-30 cm deep) of Herzliya marina (32° 09′ 38.8” N, 34° 47′ 35.0” E), Israel, during Autumn of 2019, 2020, 2021 and Spring of 2020. Before sampling, bags were washed thoroughly with filtered (0.2 µm) sterile artificial seawater (FSW). For DNA extraction, samples were kept immersed in FSW on ice until processing. For Scanning Electron Microscopy (SEM), samples were transferred to 4% PFA and 1% glutaraldehyde fixation solution for 2-5 hours and then washed with 1x PBS and kept in 1:1 ethanol/PBS solution at - 20 °C until use.

### Microscopy (LM and SEM)

For visualization of the live plastisphere flora and fauna and Lactophenol Cotton Blue (LFCB)-stained samples, we used the Nikon Eclipse Ci-L microscope with x10, x20, and x40 objectives. Photos were taken with Nikon DS-Fi3 (CMOS) digital camera. For SEM imaging, we followed our previously tested protocol (Davidov, Iankelevich-Kounio et al. 2020). In brief, fixed samples were dehydrated in graded ethanol series of 50%, 70%, 85%, and 95% ethanol (10 min each), followed by 3×15 min in 100% ethanol. Dehydrated samples were air-dried for at least 5 h in a hood, and sputter-coated with 10 nm of platinum/gold (Quorum Q150T ES). Samples were visualized and imaged using Ultra High-Resolution Maia 3 FE-SEM (Tescan) in a range of 3–7 kV voltage.

### LFCB staining

In order to identify fungal structures (hyphae, sporangia & spores), Lactophenol Cotton Blue (LFCB) dye was used. The dye is based on Methyl Blue dye with a Lactophenol solution (Phenol C_6_H_5_OH, Lactic acid CH_3_CH(OH)COOH, and Glycerol C_3_H_8_O_3_). LFCB dye stains sugar molecules including glucans and chitin, that compose the fungal cell wall (Larone and Larone 1987). Plastic samples were cut to an appropriate size for microscopy and were placed in a petri dish with 2-3 drops of LFCB for 5 mins and then washed with 1x PBS. Lastly, washed samples were placed on the top of a microscope glass slide and covered with a cover slip.

### DNA extraction and amplification of ITS barcode

For molecular identification of the symbiotic fungi, we relied on a short barcoding region within the internal transcribed spacer (ITS) locus that was previously shown to be comprehensive and specific to fungi (Schoch, Seifert et al. 2012). For DNA isolation, three plastic pieces (1mm x 1mm) with diatom-baring fungal hyphae were cut under LM and placed into a 1% Sodium Dodecyl Sulfate Buffer (SDS) buffer. Samples were then lysed in 100 µg/ml Proteinase-K in 1% SDS buffer with ceramic beads for 1 hour at 56°C while mixing thoroughly every 15 minutes. The following steps were performed using QIAamp DNA kit (Qiagen) according to the manufacturer’s instructions.

For PCR amplification, ∼10 ng of template DNA was used in 50 µl reaction volume. The PCR reactions included the sample DNA, a positive (yeast DNA), and negative (no template) controls. The ITS barcode was amplified using the ITS4 and ITS86 primers in a PCR reaction which was carried out with the following steps: 2 minutes at 94°C, 32 cycles of 30 seconds at 94°C, 30-90 seconds at 45°C to 57°C, 30-90 seconds at 72°C, and final extension at 72°C for 5 minutes. PCR product (1 µl) was run in 1% agarose gel (100 volts, 25 min) alongside a PCR Bio IV DNA size marker. The PCR product was cleaned up on magnetic rack with SPRI magnetic beads (Canvax, Spain) following the manufacturer’s instructions.

### Nanopore MinION sequencing

Because we isolated a “dirty” sample of the symbiotic fungus, ITS metabarcoding approach was used for the taxonomic identification, under the assumption that the target sequence is expected to generate the most abundant PCR product, as was previously demonstrated (Maestri, Cosentino et al. 2019). The Nanopore MinION sequencing libraries were prepared using the 1D Native barcoding genomic DNA protocol with EXP-NBD 104 and SQK-LSK 109 kits (Oxford Nanopore Technologies). ∼200 fmol of purified amplification products were subjected to DNA repair and end-prep using a NEBNext DNA repair mix and NEBNext Ultra II End Repair/dA-Tailing Module (New England Biolabs). The library preparation included two ligation steps. In the first step, the multiplexing barcodes were ligated to the amplicons, using T4 Ligase (New England Biolabs) and then equal molarities of the barcoded amplicons were pooled together. In the second step, the MinION adaptor was ligated to the amplicons of the pooled sample. Each step was followed by DNA purification with SPRI magnetic beads. ITS sequencing library was loaded to a new MinION flow cell (R9.4.1) and sequenced. Base-calling for all libraries was done automatically by the MinKNOW program using the “high accuracy” base-caller option. Raw reads were obtained in FAST5 and formats from which “pass” quality reads were subjected to further analysis.

### Data analysis

The raw nanopore reads were demultiplexed and trimmed with qcat (https://github.com/nanoporetech/qcat). Primers were removed using Cutadapt with 10% error tolerance. The reads were filtered based on read quality and read length (300-340bp), based on the plotted quality/length distribution similar to (Davidov, Iankelevich-Kounio et al. 2020). The filtered reads were used to create a single consensus sequence using the ONTrack pipeline (https://github.com/MaestSi/ONTrack) according to (Maestri, Cosentino et al. 2019). The consensus sequence was deposited in the genebank (accession no. OQ843024) and was compared with NCBI and Silva databases for taxonomic identification. In order to assess the relative abundance of the symbiotic fungi, the consensus sequence was compared to an ITS database that was created from samples that were incubated in the same marina location for similar time periods (Marsay, Koucherov et al. 2022). The comparison was conducted with local blast+ suite (Version 2.13.0) provided by the National Center for Biotechnology Information (NCBI), wherein a threshold of 80% query coverage and 94% sequence identity were applied. Prism software (v 8.0.2) was used to evaluate significant differences with Holm-Sidak’s multiple comparisons test and to create the graphs.

## Results

### Interactions of diatoms with fungal hyphae

LDPE plastic surfaces that were incubated for one month during autumn 2019, in the Herzliya marina (Israel), contained a rich biofilm that could be noticed by the naked eye. Light and electron microscope (SEM) images revealed a network of fungal hyphae with a few fruiting bodies (Fig. 1A). Some of the fungal hyphae contained diatoms attached to them. The fungi-attached diatoms were of different species belonging to several pennate diatom genera including *Striatellaceae* (Fig. 1B), *Licmophora* (Figs. 1B, 1N), *Nitzschia* (Figs. 1D, 1H, 1L), *Amphora* (Figs. 1K and 1L) and *Achnathes* (Fig. 1M). The diatom-baring hyphae were positive to lactophenol cotton blue staining (Fig 1C), confirming the presence of typical fungal cell walls. Both fungus-bound and free diatom cells were often found in couples - after a single division (Figs. 1C, 1H), or as cell bundles after multiple divisions (Figs. 1D, 1N). Dividing diatom cells were found to be attached from one of the frustule’s poles, while non-dividing individuals were mostly observed attached from their perpendicular plane. The fungus-diatom contact points were enriched by unknown extracellular material (e.g., Fig. 1H), which presumably strengthened the anchoring of the diatom cells to the hyphae filaments.

**Fig. 1.**
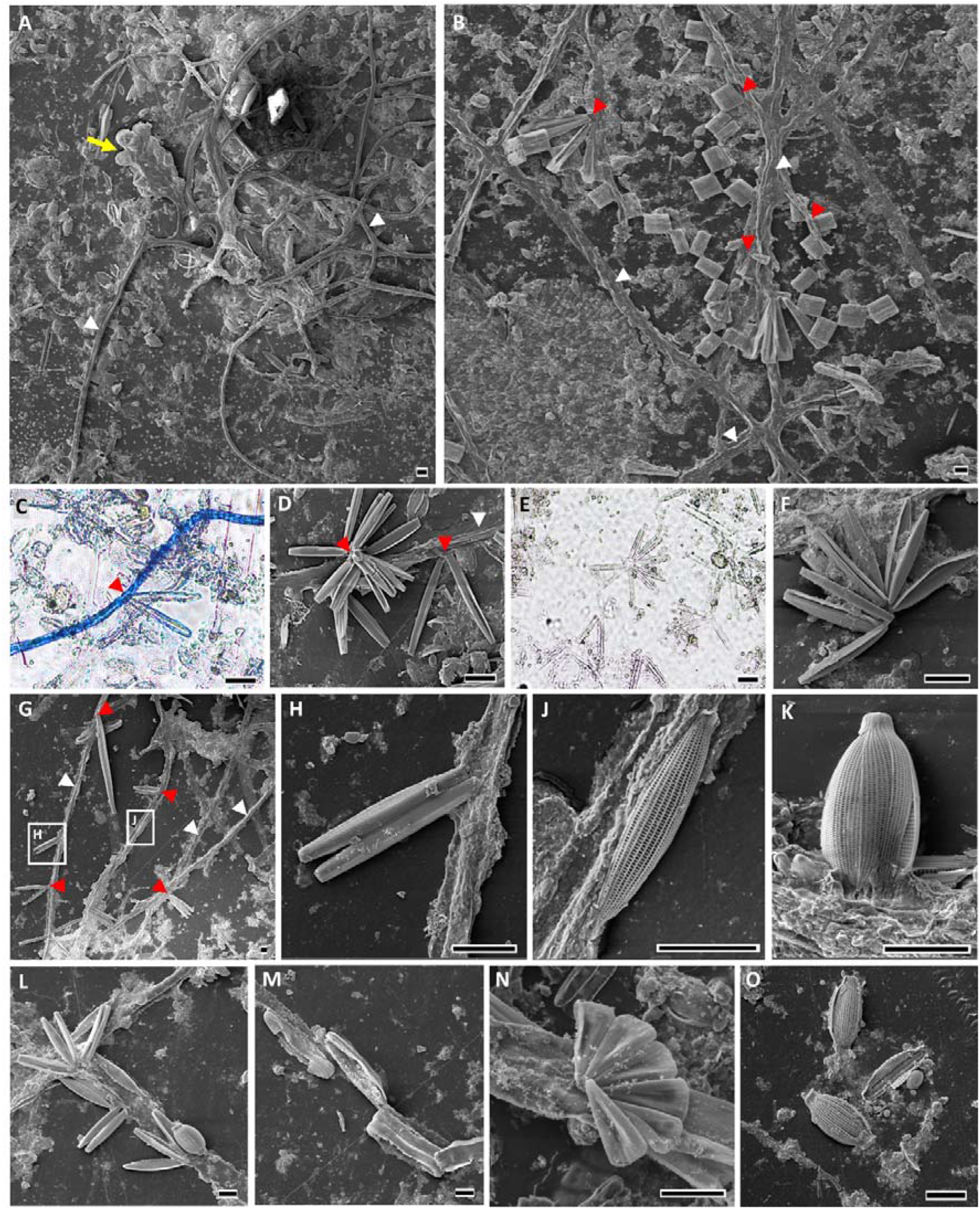
Symbiotic relations between fungi and diatoms on LDPE surface. Pictures C and E were taken using LM. All other figures were taken using SEM. A. a network of fungal hyphae and a fruiting body (yellow arrow) attached to the surface. B. a section of the surface covered with a network of fungal hyphae and diatoms from the genera *Striatellaceae* (rectangular) and *Licmophora* (elongated) attached to them. C. fungal hypha stained with lactophenol cotton blue with unstained post-division diatoms attached to it. D. post-division *Nitzschia* sp. attached to fungal hypha. E, F. the same diatom species as in D attached directly to the plastic surface. G. a section of the surface covered with a network of fungal hyphae and diatoms attached to them. H and J. enlargements of elongated diatoms attached to fungal hypha in different positions K. *Amphora* sp. frustule attached to hypha partially covered with secreted fungal substance. L-N. different diatom genera on fungal hyphae including *Nitzschia* and *Amphora* (L), *Achnanthes* (M), and *Licmophora* (N). O. Amphora diatoms on the plastic surface. White arrowheads indicate fungal hyphae. Red arrowheads indicate fungal-diatom contact points. Scale: 10µm

### The nature of the fungal-algal relationships

As fungal-diatom firm attachment was repeatedly observed in several LDPE surfaces, the relationships were referred to as symbiotic relationships. To define whether the symbiosis is obligatory or facultative, we searched for diatoms of the same species that are not attached to fungi elsewhere on the plastic surface. Indeed, unassociated diatom cells (both single and post-division) of the same species were found, indicating that the symbiosis is facultative. To understand the prevalence of the phenomenon we calculated the ratio between bound vs. unbound diatoms in three 1x1 cm surfaces. The most common fungus-bound diatom of genus *Nitzschia* (Fig. 1D, 1L) had an almost equal number of bound vs. unbound cells (∼48%). We note that the observed symbiosis was not recorded in other LDPE incubation experiments that were conducted during spring 2020 but were found again during autumn 2020 and autumn 2021.

### Diatom-baring hyphae belong to an unknown Ascomycota fungus with a preference for PE surfaces

While diatoms may often be classified using high-resolution microscopy images up to the genus and even to the species level, this is not the case for marine fungi. We therefore implemented the DNA barcoding approach to classify the taxon of the symbiotic fungus based on a fraction of the ITS locus (ITS4 - ITS86). Although DNA was extracted from a small piece of fungus-containing surface, it is reasonable to assume that it was contaminated with other fungal DNA traces. This notion was supported by the large PCR amplicon size range which was observed on the gel image (Fig 2A) and in the analysis of the amplicon read sizes (Fig 1B). We therefore used nanopore MinION high throughput sequencing to identify the most prevalent and repetitive ITS barcode sequence. To reduce background noise, analysis was performed on a restricted sequence length range based on the amplicon size distribution (Fig. 2B). The results showed a single consensus of 284 bp (Acc. OQ843024) which was mapped to an unknown taxon from phylum Ascomycota (referred to as fungus “F1”). We further used the consensus sequence to screen for similar sequences in our previously published ITS database from the same marine environment (Marsay, Koucherov et al. 2022). The results showed with high probability that F1 fungus was generally more abundant on surfaces vs. water with a significant preference for LDPE surfaces over polyethylene terephthalate (PET) and glass surfaces (Fig. 2C).

**Fig. 2.**
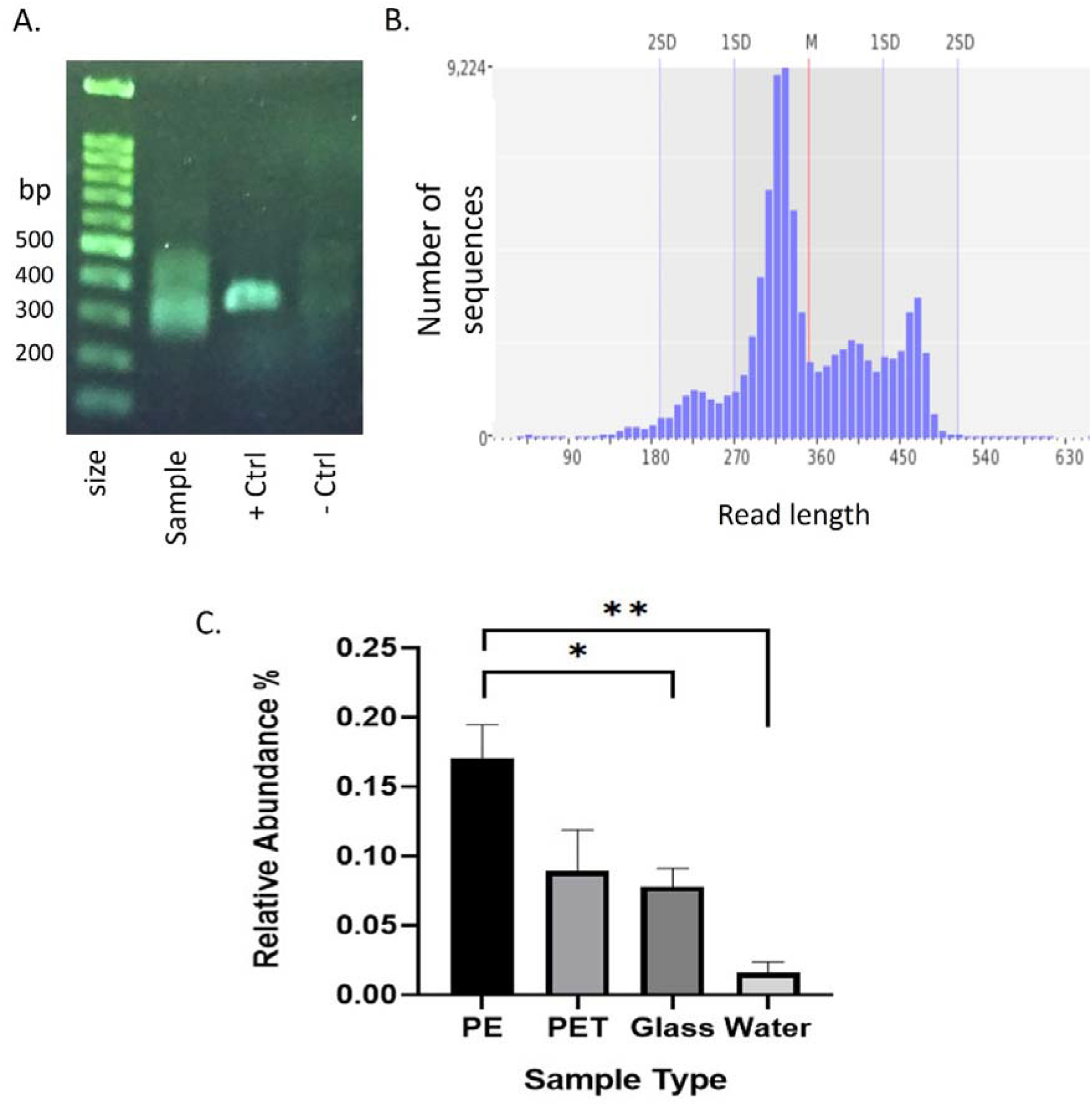
Molecular barcoding and plastic surface preference of an unknown diatom-baring F1 fungus. A, B. The ITS barcode amplification of the fungal hyphae by PCR resulted in amplicons within a large range of sizes peaking at 320±20 bp. C. relative abundance of F1 fungus on different surfaces vs. water based on the comparison of its ITS 284 bp-long consensus to the previously published ITS database (Marsay, Koucherov et al. 2022) with a similarity threshold of 94%.

## Discussion

Terrestrial mutualistic relationships between plants and fungi are very common in nature. Bioinformatic analyses suggested that the molecular basis for the plant-fungi symbiosis originated before land colonization by plants and that it preexisted in their algal ancestors (Delaux, Radhakrishnan et al. 2015). In lichens, fungi and unicellular algae reside together in mutualistic and often obligatory symbiotic relationships. The unique partnership within lichens allows life in extreme habitats which neither fungi nor algae alone can survive (Fernández□Mendoza, Domaschke et al. 2011). The photobionts within lichens contribute metabolically by secretion of sugars that are absorbed by fungi. In return, the fungi provide the photobiont (either algae or cyanobacteria) with a moist habitat, protection from UV, and a steady CO_2_ supply (Nash 1996). Lichenized fungi are now commonly referred to a separate group within the fungi kingdom (Grube and Wedin 2016).

In this study, we characterize a novel symbiotic relationship between fungi and several species of pennate diatoms on a plastic surface that was submerged in the water of a Mediterranean Sea marina for a period of one month. In a previous study, we identified a network of fungal hyphae that partially covered plastic surfaces that were submerged in the marina. As we did not identify a similar network on plastic debris that was collected in the open sea, we assume that the relatively undisturbed, yet polluted (with organic waste) marina water contribute to the development of such networks. The symbiosis was observed in the autumns of three consecutive years (2019-2021) but was not found in spring. This seasonality is not surprising given the overall taxonomic seasonality observed within the plastic biome (Marsay, Ambrosino et al. 2023). Furthermore, according to our ITS barcode analysis and the comparisons to our previously established database, we concluded that the symbiotic fungus is more prevalent on PE surfaces vs. PET or glass surfaces. PE is generally less dense than seawater, therefore tends to float near the water surface, while PET and glass sinks. Thus, the observed preference may reflect an adaption for colonizing surfaces that allow photosynthesis by the photobiont (the diatoms). While we did not investigate the mechanism of the symbiosis, we suggest that both the fungus and the diatoms gain benefits from it. We suggest that the diatoms use the fungal hyphae as an anchoring network to the floating surface. This notion is supported by the unknown secreted fungal substance, observed by SEM, which seems to glue the diatoms to the hyphae filaments. This anchoring may allow frequent diatom cell division without losing grip of the surface as was often observed. On the other hand, similar to lichens, the fungus may utilize sugars secreted by the diatoms or even diatom corps (that were observed attached to hyphae).

As marine plastic surfaces are relatively new in our world, it is possible that the observed plastic-associated symbiosis was adapted from a preexisting symbiosis that occurs in either shallow benthic marine environments, natural floating organic surfaces such as wood and leaves, or even on living organisms such as multicellular algae. Further research may reveal ancestral symbiosis genes that in turn may promote our understanding of fungal-algal symbiotic relationships.

## References

Amaral-Zettler, L. A., E. R. Zettler and T. J. Mincer (2020). “Ecology of the plastisphere.” Nature Reviews Microbiology 18(3): 139–151.

Amaral□Zettler, L. A. (2022). “Colonization of plastic marine debris: The known, the unknown, and the unknowable.” Plastics and the Ocean: Origin, Characterization, Fate, and Impacts: 301–316.

Armbrust, E. V. (2009). “The life of diatoms in the world’s oceans.” Nature 459(7244): 185–192.

Benoiston, A.-S., F. M. Ibarbalz, L. Bittner, L. Guidi, O. Jahn, S. Dutkiewicz and C. Bowler (2017). “The evolution of diatoms and their biogeochemical functions.” Philosophical Transactions of the Royal Society B: Biological Sciences 372(1728): 20160397.

Carson, H. S., M. S. Nerheim, K. A. Carroll and M. Eriksen (2013). “The plastic-associated microorganisms of the North Pacific Gyre.” Marine pollution bulletin 75(1-2): 126–132.

Davidov, K., E. Iankelevich-Kounio, I. Yakovenko, Y. Koucherov, M. Rubin-Blum and M. Oren (2020). Identification of plastic-associated species in the Mediterranean Sea using DNA metabarcoding with Nanopore MinION. Sci Rep 10: 17533.

Delaux, P.-M., G. V. Radhakrishnan, D. Jayaraman, J. Cheema, M. Malbreil, J. D. Volkening, H. Sekimoto, T. Nishiyama, M. Melkonian and L. Pokorny (2015). “Algal ancestor of land plants was preadapted for symbiosis.” Proceedings of the National Academy of Sciences 112(43): 13390–13395.

Du, Y., X. Liu, X. Dong and Z. Yin (2022). “A review on marine plastisphere: biodiversity, formation, and role in degradation.” Computational and Structural Biotechnology Journal.

Eriksen, M., W. Cowger, L. M. Erdle, S. Coffin, P. Villarrubia-Gómez, C. J. Moore, E. J. Carpenter, R. H. Day, M. Thiel and C. Wilcox (2023). “A growing plastic smog, now estimated to be over 170 trillion plastic particles afloat in the world’s oceans—Urgent solutions required.” Plos one 18(3): e0281596.

Fernández□Mendoza, F., S. Domaschke, M. García, P. Jordan, M. P. Martín and C. Printzen (2011). “Population structure of mycobionts and photobionts of the widespread lichen Cetraria aculeata.” Molecular Ecology 20(6): 1208–1232.

Gkoutselis, G., S. Rohrbach, J. Harjes, M. Obst, A. Brachmann, M. A. Horn and G. Rambold (2021). “Microplastics accumulate fungal pathogens in terrestrial ecosystems.” Scientific Reports 11(1): 13214.

Grube, M. and M. Wedin (2016). “Lichenized fungi and the evolution of symbiotic organization.” Microbiology spectrum 4(6): 4.6. 34.

Kim, S. H., J. W. Lee, J. S. Kim, W. Lee, M. S. Park and Y. W. Lim (2022). “Plastic-inhabiting fungi in marine environments and PCL degradation activity.” Antonie Van Leeuwenhoek 115(12): 1379–1392.

Lacerda, A. L. d. F., M. C. Proietti, E. R. Secchi and J. D. Taylor (2020). “Diverse groups of fungi are associated with plastics in the surface waters of the Western South Atlantic and the Antarctic Peninsula.” Molecular Ecology 29(10): 1903–1918.

Larone, D. H. and D. H. Larone (1987). Medically important fungi: a guide to identification, Citeseer.

Maestri, S., E. Cosentino, M. Paterno, H. Freitag, J. M. Garces, L. Marcolungo, M. Alfano, I. Njunjić, M. Schilthuizen and F. Slik (2019). “A rapid and accurate MinION-based workflow for tracking species biodiversity in the field.” Genes 10(6): 468.

Marsay, K. S., A. C. Ambrosino, Y. Koucherov, K. Davidov, N. Figueiredo, I. Yakovenko, S. Itzahri, M. Martins, P. Sobral and M. Oren (2023). “The geographical and seasonal effects on the composition of marine microplastic and its microbial communities: The case study of Israel and Portugal.” Frontiers in Microbiology 14.

Marsay, K. S., Y. Koucherov, K. Davidov, E. Iankelevich-Kounio, S. Itzahri, M. Salmon-Divon and M. Oren (2022). “High-Resolution Screening for Marine Prokaryotes and Eukaryotes With Selective Preference for Polyethylene and Polyethylene Terephthalate Surfaces.” Frontiers in Microbiology: 1166.

Nash, T. H. (1996). Lichen biology, Cambridge University Press.

Philippe, A., C. Noël, B. Eyheraguibel, J.-F. Briand, I. Paul-Pont, J.-F. Ghiglione, E. Coton and G. Burgaud (2023). “Fungal Diversity and Dynamics during Long-Term Immersion of Conventional and Biodegradable Plastics in the Marine Environment.” Diversity 15(4): 579.

Reisser, J., J. Shaw, G. Hallegraeff, M. Proietti, D. K. Barnes, M. Thums, C. Wilcox, B. D. Hardesty and C. Pattiaratchi (2014). “Millimeter-sized marine plastics: a new pelagic habitat for microorganisms and invertebrates.” PloS one 9(6): e100289.

Round, F. E., R. M. Crawford and D. G. Mann (1990). Diatoms: biology and morphology of the genera, Cambridge university press.

Schoch, C. L., K. A. Seifert, S. Huhndorf, V. Robert, J. L. Spouge, C. A. Levesque, W. Chen, F. B. Consortium, F. B. C. A. List and E. Bolchacova (2012). “Nuclear ribosomal internal transcribed spacer (ITS) region as a universal DNA barcode marker for Fungi.” Proceedings of the national academy of Sciences 109(16): 6241–6246.

Vyverman, W. (2004). “Experimental studies on sexual reproduction in diatoms.” Int. Rev. Cytol 237: 91.

Zettler, E. R., T. J. Mincer and L. A. Amaral-Zettler (2013). “Life in the “plastisphere”: microbial communities on plastic marine debris.” Environmental science & technology 47(13): 7137–7146.

